# Evaluating proteomics imputation methods with improved criteria

**DOI:** 10.1101/2023.04.07.535980

**Authors:** Lincoln Harris, William E. Fondrie, Sewoong Oh, William S. Noble

## Abstract

Quantitative measurements produced by tandem mass spectrometry proteomics experiments typically contain a large proportion of missing values. This missingness hinders reproducibility, reduces statistical power, and makes it difficult to compare across samples or experiments. Although many methods exist for imputing missing values in proteomics data, in practice, the most commonly used methods are among the worst performing. Furthermore, previous benchmarking studies have focused on relatively simple measurements of error, such as the mean-squared error between the imputed and the held-out observed values. Here we evaluate the performance of a set of commonly used imputation methods using three practical, “downstream-centric” criteria, which measure the ability of imputation methods to reconstruct differentially expressed peptides, identify new quantitative peptides, and improve peptide lower limit of quantification. Our evaluation spans several experiment types and acquisition strategies, including datadependent and data-independent acquisition. We find that imputation does not necessarily improve the ability to identify differentially expressed peptides, but that it can identify new quantitative peptides and improve peptide lower limit of quantification. We find that MissForest is generally the best performing method per our downstream-centric criteria. We also argue that exisiting imputation methods do not properly account for the variance of peptide quantifications and highlight the need for methods that do.

## 1 Introduction

The quantitative accuracy and sensitivity of tandem mass spectrometry proteomics experiments has increased dramatically in the past decade. In spite of this trend, proteomics experiments are still limited by excessive “missingness,” which refers to peptides that are present in the sample matrix but are not detected by the instrument. Missingness can be attributed to a variety of technical factors including ion suppression, coeluting peptides, the lower limit of quantification of the instrument, and the failure to confidently assign peptides to all observed spectra [1, 2]. Although low abundance peptides are generally more likely to be missing, peptides may be missing across the entire range of intensities. Missingness decreases the statistical power of proteomics experiments, hinders reproducibility, and makes it difficult to compare across batches or experiments [1, 2].

Imputation is a bioinformatic solution to the missingness problem. Imputation entails using statistical or machine learning procedures to estimate missing values in a data set. While still not widely accepted in the proteomics community, imputation has been standard practice for decades for analysis of gene expression [3] and clinical and epidemiological data [4], and more recently astronomy [5, 6] and single-cell transcriptomic data [7, 8]. Imputation methods for proteomics data (Table 1) fall into three broad categories: “single-value replacement” methods, in which all missing values are filled in with a single replacement value; “local similarity” methods, which use statistical models to learn patterns of local similarity in the data, for example between subsets of similar peptides or runs; and “global similarity” methods, which learn broad patterns of similarity across all peptides and runs.

**Table 1.** Existing proteomics imputation methods. Descriptions of general categories of imputation strategies and examples of specific tools that implement each strategy. The “Type” column indicates whether the method uses single-value replacement (S), local similarity (L), or global similarity (G).

| Method | Type | Description | Examples |
| --- | --- | --- | --- |
| Zero replacement | S | Replace missing values with zeros |  |
| Mean replacement | S | Replace missing values with the mean peptide intensity for a peptide or sample |  |
| Low value replacement | S | Replace missing values with the lowest observed intensity in any sample (sample minimum) or peptide (peptide minimum) |  |
| Gaussian random sample | S | Randomly sample from a Gaussian distribution centered around the lowest observed intensity | Perseus [9] |
| Regression | L | Linear regression is used to estimate missing values | lm, glm |
| kNN | L | Weighted average intensity of $k$ most similar peptides | impute.knn [3], VIM [10] |
| MissForest | G | Nonparametric method to impute missing values using a random forest classifier trained on the observed parts of the data set, repeated until convergence | MissForest [11] |
| PCA | G | Run principal component analysis, impute missing values with the regularized reconstruction formulas and repeat until convergence | pcaMethods [12], missMDA [13] |

It is not always clear what imputation method is best for a given proteomics data set. A number of studies benchmark imputation methods and offer guidelines for selecting an appropriate method [1, 2, 14–17]. A general recommendation is that single-value replacement strategies rarely work well. Another is that the optimal imputation method depends on the structure of missingness in the data. Mass spectrometry-based proteomics experiments exhibit two major forms of missingness: missing completely at random (MCAR) and missing not at random (MNAR). MCAR describes the case in which missingness does not depend on any observed variable, that is, missingness occurs independent of peptide intensity or relationships between samples. In the case of MNAR, missingness *is* dependent on some observed variable. For example, in mass spectrometry-based proteomics, missingness is often a function of peptide intensity, with more missingness occurring in peptides closer to the instrument’s lower limit of quantification (LLOQ).

Most proteomics imputation method benchmarking studies use relatively simple performance measures that are not necessarily relevant to proteomics researchers. One common example is computing the mean squared error (MSE) between imputed and ground truth peptides quantifications for a withheld set of matrix entries. We argue that such evaluation metrics, while certainly valid, are neither the most relevant nor the most useful to proteomics researchers. We therefore introduce alternative, “downstream-centric,” criteria focused on differential expression, peptide LLOQ, and the total number of quantitative peptides in an experiment. We argue that these downstream-centric criteria are more relevant to the questions proteomics researchers typically seek to answer. Furthermore, we observe that the best-performing imputation methods per traditional criteria often differ from the best performing methods per our downstream-centric criteria.

To decide which imputation methods to include in our study, we carried out a systematic literature review. All *Journal of Proteome Research* articles published between January 1, 2019, and January 31, 2023, were searched for the following terms: “impute,” “imputed,” “imputation.” For this survey we excluded methodological and benchmarking studies. On the basis of the resulting citation counts (Figure 1), we selected four of the most popular imputation methods: *k*-nearest neighbor (kNN) [3], MissForest [11], Gaussian sampling [9], and low value replacement (Figure 1). We also include a non-negative matrix factorization (NMF) imputation method, which has recently been proposed for proteomics [18–20]. By focusing only on the most commonly used imputation methods, our aim is to provide a practical comparison that will be beneficial to experimental proteomicists. For this reason, seldom used R packages (e.g., imp4p, impSeqRob, QRLIC) have been omitted from our analysis. We also omit PCA-based methods, as we did not find a single recent study that used them. Additionally, we choose to conduct our analysis exclusively on peptide-level quantifications. Our reason for this is twofold: (i) summarizing peptide quantifications at the protein level reduces often-critical data heterogeneity [21], and (ii) imputation generally performs better at the peptide level [15].

**Figure 1.**
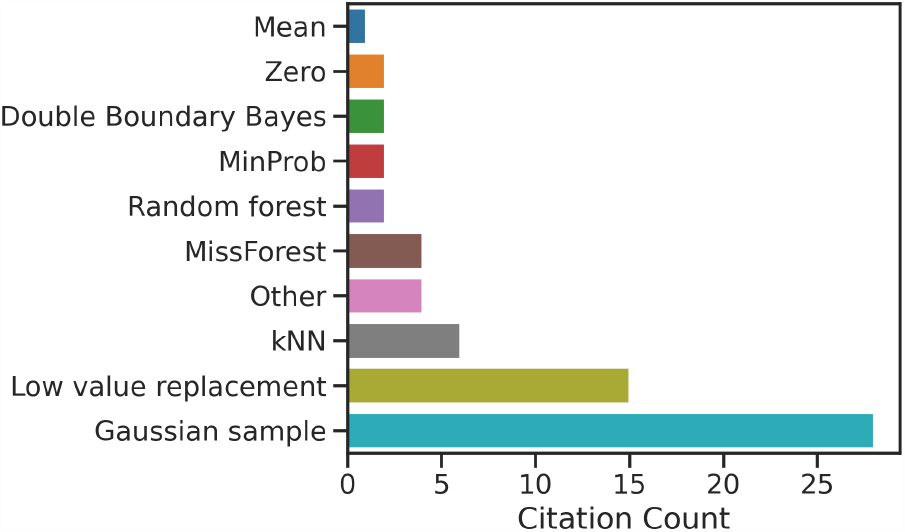
Citation counts for the most commonly used proteomics imputation methods. Results of a literature survey of *Journal of Proteome Research* articles from January, 2019 – January, 2023 are shown. Methods labeled “other” appear in just a single publication, and refer to the imp4p and QRLIC R packages, as well as methods based on Euclidean distances and randomly drawing from the entire peptide intensity range. The full results of this literature search, including the names and DOIs of the identified studies, are included as Supplementary File 1.

We evaluate the performance of each of the five selected methods with both traditional and downstream-centric criteria. The latter include the ability of an imputation method to: (i) reconstruct differentially expressed peptides, (ii) identify new quantitative peptides, and (iii) improve the LLOQ of peptides in a proteomics experiment. Our benchmarking study consists of a variety of data set types, including a serial dilution experiment, data-dependent and data-independent acquisition (DDA and DIA) studies, as well as label-free and isobaric labeled experiments. Critically, we include an unimputed condition for all three downstream-centric evaluation experiments, for evaluating whether imputation should be performed in the first place. Our findings suggest that imputation may not significantly improve the ability to detect differentially expressed peptides, but that it can identify new quantitative peptides and improve peptide LLOQ.

We also demonstrate that peptide quantifications exhibit greater than expected variance in relation to measured intensity. Ion detection is a Poisson process, and so quantifications from ion-counting mass spectrometers are often assumed to be Poisson distributed [22, 23]. We demonstrate that peptide quantification measurements are overdispersed and are thus not Poisson distributed, and that log transformation does not completely correct for this overdispersion. This finding is important because many existing imputation methods make distributional assumptions that are not actually met by mass spectrometry-based proteomics data. To our knowledge, no existing method models quantitative proteomics data with the proper distributional assumptions. This suggests the need for methods that employ variance stabilization prior to imputation, similar to strategies taken in genomics [24–26].

## 2 Methods

### 2.1 Data sets

For this study, we used 12 public quantitative proteomics data sets (Table 2). Nine of the 12 data sets were accessed via the Proteomics Identification Database (PRIDE, https://www.ebi.ac.uk/pride/) [27], and are indicated with their ProteomeXchange (PXD) labels [28]. The remaining two data sets were obtained from the National Cancer Institute’s Clinical Proteomic Tumor Analysis Consortium (CPTAC) data portal (https://proteomic.datacommons.cancer.gov/pdc/) [29]. A complete list of the files obtained for each experiment is provided in Supplementary File 2.

**Table 2.**
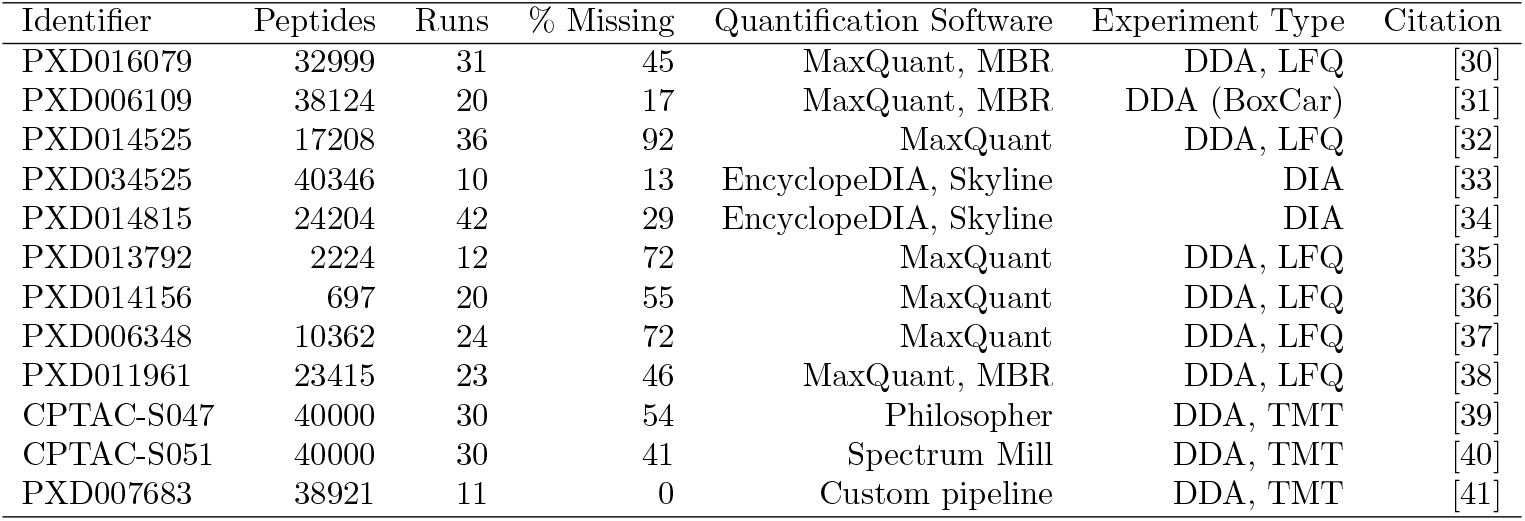
Data set characteristics. Description of the public proteomics data sets used in this study. The two CPTAC data sets were downsampled by randomly selecting 40,000 peptides and 30 runs each. “MBR” stands for “match between runs,” “LFQ” for “label-free quantification,” “SPS” for “synchronous precursor selection”, and “TMT” for “tandem mass tag.” Quantification software references: MaxQuant [42], EncyclopeDIA [43], Skyline [44], Philosopher [45].

For experiments processed with MaxQuant, we used the provided “peptides.txt” files to generate peptideby-run intensity matrices by selecting only the “Sequence” and “Intensity” columns. For experiments not processed with MaxQuant, we obtained peptide-spectrum match files and converted them to matrix format with custom scripts (available at https://github.com/Noble-Lab/2023-prot-impute-benchmark). The peptide quantification matrices from the two CPTAC studies were large (S047: 110k peptides *×* 226 samples; S051: 291k peptides *×* 35 samples). For efficiency, we downsampled both matrices by randomly selecting 40,000 peptides and 30 runs from each.

### 2.2 Traditional evaluation measures

We first used a traditional train/test setup to evaluate the performance of imputation methods. In this approach, the values in the matrix are randomly segregated into two groups: a training set and a test set. The imputation method is given the training set values, and we measure how well the method imputes the values in the test set. For each data set, peptides with fewer than four present values in the training set were removed prior to splitting. These tests were carried out using the following seven data sets: PXD014156, PXD011961, PXD014525, CPTAC-S051, CPTAC-S047, PXD034525, PXD014815 (Table 2).

For each data set, we simulated missingness according to two procedures: MCAR and MNAR. For MCAR, 25% of the present (i.e., non-missing) matrix entries were randomly selected for the test set. The remaining matrix entries were used as the training set. For MNAR, we took a similar approach to the one described by Lazar *et al*. [15]. For a given peptide quantifications matrix, we constructed an equally sized *thresholds matrix* filled with values sampled from a Gaussian distribution centered about the 30^th^ percentile of the distribution of quantifications, with standard deviation 0.6. For each element *X*_*ij*_ in the peptides matrix, if the corresponding thresholds matrix element *T*_*ij*_ *< X*_*ij*_, then *X*_*ij*_ was assigned to the training set. Otherwise, a single Bernoulli trial with probability of success 0.75 was conducted. If this Bernoulli trial was successful, then *X*_*ij*_ was assigned to the test set. Otherwise *X*_*ij*_ was assigned to the training set. Bernoulli success probability and Gaussian distribution mean and standard deviation were selected in such a way that 25% of present matrix entries were ultimately assigned to the test set. The remaining 75% were assigned to the training set. The distributions of the training and test set values following the MCAR and MNAR partitions are shown in Supplementary Figure S1, for experiment PXD034525.

Once missing values had been simulated into the six matrices, imputation was performed with five procedures: NMF, kNN, MissForest, low value replacement (sample minimum) and Gaussian sample impute. A custom PyTorch model was used for NMF imputation. This model used an MSE loss function and stochastic gradient descent to converge on an ideal matrix factorization. This model is available at https://github.com/Noble-Lab/ms_imputer. For kNN, we used the KNNImputer implementation from scikit-learn. MissForest version 1.5 was used (https://CRAN.R-project.org/package=missForest) [11]. Custom code was used for the low value replacement and Gaussian sample impute procedures. For Gaussian sample impute we attempted to replicate the procedure taken by Perseus [9]. For low value replacement, we filled in missing values with the lowest measured peptide intensity for each sample. NMF and kNN were performed with four latent factors and neighbors, respectively. MissForest was performed with 100 trees, the default setting.

Following imputation, we computed the MSE between observed and imputed values for each test set.

### 2.3 Downstream-centric evaluation measures

#### 2.3.1 Differential expression

For differential expression analysis we obtained data from PXD034525, a DIA study of Alzheimer’s disease [33]. Clinical samples had previously been assigned to experimental groups based on several genetic, histopathological and cognitive criteria. We compared differentially expressed peptides between (i) autosomal dominant Alzheimer’s disease dementia and (ii) high cognitive function, low Alzheimer’s disease neuropathologic change. Both experimental groups were composed of nine samples—each obtained from a separate patient—and 32,614 detected peptides.

Ground truth differentially expressed peptides were determined by performing two-sample t-tests between experimental groups for each detected peptide. P-values were corrected for multiple hypothesis testing using the Benjamini-Hochberg procedure [46]. Peptides with corrected p-values *<* 0.01 were considered ground truth differentially expressed.

MCAR and MNAR partitioning was performed similar to above, but this time we created three disjoint sets: training, validation and test. For the MCAR partition, 15% of matrix entries were randomly selected without replacement for the validation set, and a separate 15% were selected for the test set. For the MNAR partition, matrix entries corresponding to successful Bernoulli trials were assigned in an alternating fashion to either the validation or the test set. Bernoulli success probability and Gaussian distribution mean and standard deviation were tuned such that 30% of present matrix entries were withheld from the training set.

The validation sets were used to select the optimal hyperparameters for NMF and kNN. For MissForest a full hyperparameter search proved computationally unfeasible, so we again selected the default value of 100 for the *n* trees parameter. None of the other methods had tunable hyperparameters. The following values were included in our hyperparameter searches for *n* latent factors and *k* neighbors: [1, 2, 4, 8, 16, 32].

Following hyperparameter selection, imputation was performed with each method. Differentially expressed peptides were determined for the imputed matrices as previously described. Precision-recall curves comparing ground truth to imputed differentially expressed peptides were generated with scikit-learn. For the unimputed condition, the differential expression calculation was performed as previous, while simply ignoring the missing matrix entries. That is, the differential expression test was performed on training set values only.

#### 2.3.2 Quantitative peptides

To examining the effects of imputation on the number of quantitative peptides in an experiment, we obtained data from PXD014815 [34]. This was a serial dilution experiment in which yeast peptides were spiked into a background matrix at various known concentrations. The authors then used a custom statistical model to fit the relationship between observed and expected signal, and to determine whether increases in signal corresponded to proportional increases in peptide abundance. Peptides in which increase in signal did indeed correspond to increases in quantity across a linear range were considered quantitative.

We used this model to assess the number of quantitative peptides before and after imputation of the serial dilutions data set with various methods. MCAR partitioning was performed as described above. Hyperparameter tuning for kNN and NMF was performed as described above. The validation and test sets were imputed with each method and quantitative peptides were identified in the imputed matrices. The UpsetR package was used to generate Figure 4 [47].

#### 2.3.3 Lower limit of quantification

We used the serial dilution experiment from PXD014815 to examine the effects of imputation on peptide LLOQ. We used the statistical model from Pino *et al*. [34] to determine the LLOQ of each detected peptide before and after imputation. One-sided binomial tests were performed to determine whether each imputation method decreases the LLOQ for significantly more peptides than it increases. Binomial p-values were corrected with the Benjamini-Hochberg procedure.

#### 2.4 Runtime evaluation

We used Python’s time module to compare runtimes of the various imputation methods. NMF, kNN, low value replacement, Gaussian sample and MissForest were run on 14 public proteomics data sets accessed from PRIDE. This experiment was performed on a dual CPU Intel Xeon E5-2620 machine with 32 GB RAM running CentrOS 7.6. NMF was specified to run on a maximum of 10 cores, and the remaining methods were run on a single core. This was because the kNN implementation we used, scikit-learn’s KNNImputer, does not support multiprocessing, nor do our custom implementations of low value replacement and Gaussian sample impute. MissForest does support multiprocessing, though in our experience, the parallelized version of MissForest proved nearly impossible to run to completion. Thus, we choose to limit MissForest to a single core.

## 3 Results

### 3.1 Evaluating with traditional criteria

We began by assessing the performance of popular imputation methods with a traditional machine learning criterion: reconstruction error on a withheld test set. Accordingly, we obtained peptide-level quantifications for seven of the experiments shown in Table 2. These include DIA, DDA and tandem mass tag (TMT) experiments, with a range of missingness from 0 to 92%. We assessed the ability of the imputation methods to reconstruct missing values, after MCAR and MNAR procedures were used to simulate an additional 25% missingness in each data set.

Our results (Figure 2) demonstrate that the best performing imputation method generally depends on the type of missingness. MissForest and NMF perform the best for all seven data sets in the MCAR condition. In the MNAR condition, the two single-value imputation methods—Gaussian sample and low value replacement—appear to work the best, though MissForest also performs well for some data sets. In both conditions the two TMT data sets, CPTAC-S047 and CPTAC-S051, yield lower reconstruction errors

**Figure 2.**
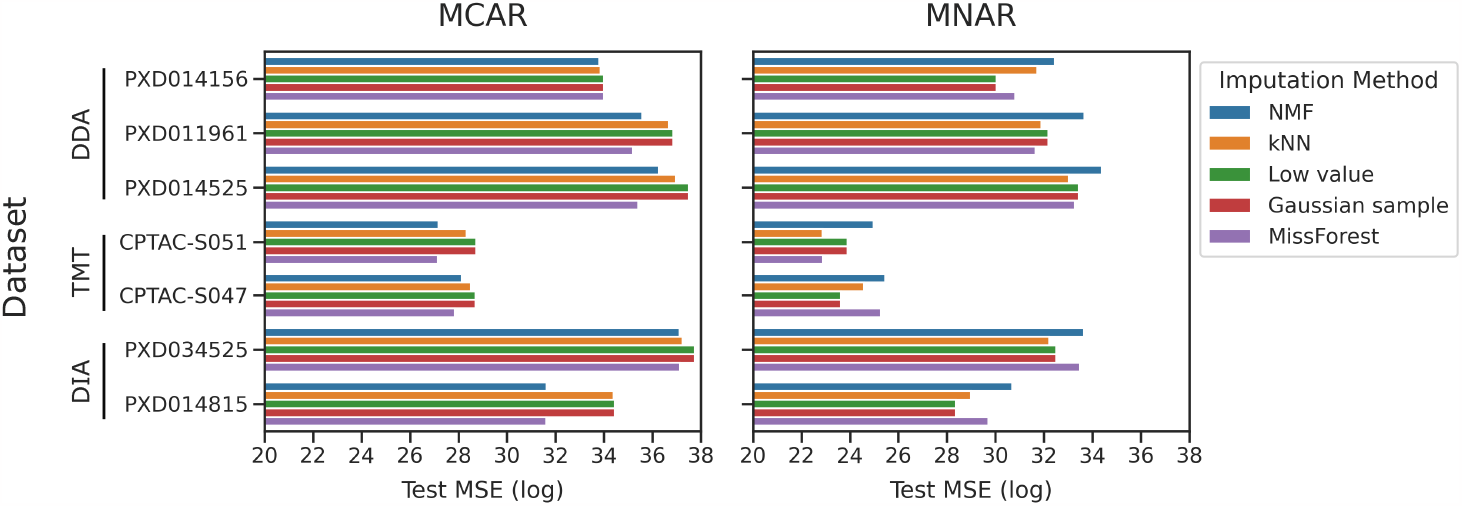
Evaluating imputation methods with traditional criteria. Reconstruction error (MSE) after imputation with five methods is shown for seven public proteomics data sets. MCAR and MNAR procedures were used to simulated missing values.

### 3.2 Evaluating with downstream-centric criteria

Although the type of evaluation shown in Figure 2 is informative, we argue that reconstruction error of a heldout set is neither the most convincing nor the most relevant evaluation metric for most proteomics researchers. Additionally, good performance on reconstruction of held-out values does not guarantee good performance on certain downstream analysis tasks. Furthermore, many existing benchmarking studies make the assumption that imputation will improve performance on some downstream analysis task relative to no imputation. This assumption is generally unfounded, as imputation can introduce bias in even the best circumstances [48, 49]. With these considerations in mind, we compared the performance of five imputation methods on three downstream analysis tasks that we argue are more congruent with the questions proteomicists typically seek to answer.

We began with differential expression analysis. We obtained peptide-level quantifications from a DIA-based clinical study of Alzheimer’s disease [33]. Merrihew *et al*. obtained brain samples and assigned them to experimental groups based on several genetic, histopathological and cognitive criteria. We compared samples belonging to two experimental groups: (i) autosomal dominant dementia and (ii) high cognitive function, low Alzheimer’s disease neuropathologic change. These experimental groups represent opposite ends of the spectrum of Alzheimer’s disease severity; thus, we reasoned that they should display significant biological heterogeneity.

We compared the abilities of the imputation methods to reconstruct peptides that are differentially expressed between the two experimental groups, in both MCAR and MNAR conditions (Figure 3). To perform this experiment, we identified ground truth differentially expressed peptides in the low-missingness DIA data set, simulated 30% missingness, then imputed with various methods and identified differentially expressed peptides in the imputed matrices. We also included an unimputed condition in which differentially expressed peptides were identified directly from the unimputed training set. Thus, the sharp elbows in the MNAR precision-recall curves are due to the fact that an alpha value of 0.01 was used for determining both ground truth and imputed differentially expressed peptides. It is likely that many peptides have corrected p-values very close to the 0.01 threshold but are not considered differentially expressed, resulting in sharp decreases in precision as soon as this threshold is crossed. Nevertheless, the trends observed in Figure 3 remain clearly interpretable.

**Figure 3.**
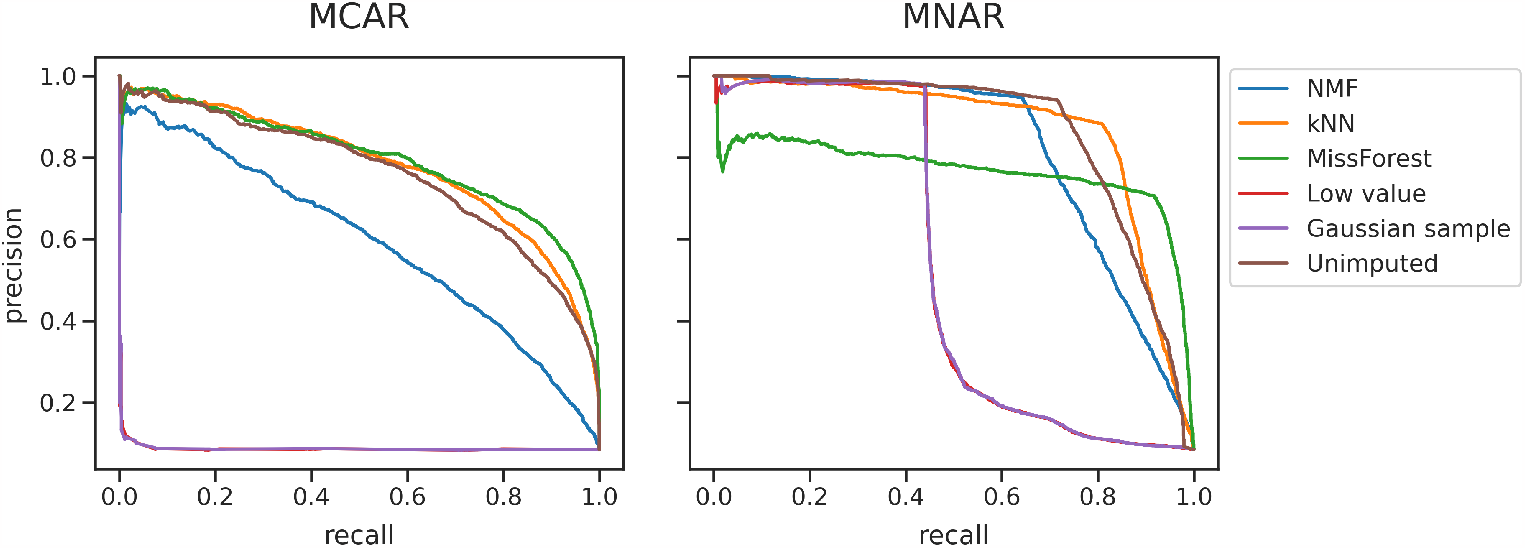
Evaluating the ability of imputation methods to reconstruct differentially expressed peptides. Precision-recall curves are shown, for MCAR and MNAR simulated missingness. Data was obtained from PXD034525, a DIA study of Alzheimer’s disease, and differentially expressed peptides were identified between high and low Alzheimer’s associated dementia groups [33]. The areas under the precisionrecall curves (AUCs) for MCAR are NMF: 0.59, kNN: 0.78, MissForest: 0.8, low value replacement: 0.09, Gaussian sample: 0.09, unimputed: 0.76. For MNAR, the AUCs are NMF: 0.82, kNN: 0.87, MissForest: 0.77, low value replacement: 0.53, Gaussian sample: 0.53, unimputed: 0.86. across all imputation methods, compared to the DDA and DIA data sets.

In the MCAR condition, MissForest, kNN and unimputed all perform well, with areas under the curve (AUCs) of 0.80, 0.78 and 0.76, respectively. In the MNAR condition, kNN, unimputed and NMF perform the best, with respective AUCs of 0.87, 0.86 and 0.82. While the two single-value imputation methods performed well in the MNAR condition of the traditional evaluation experiment (Figure 2), they perform extremely poorly on the differential expression test, with the lowest AUCs for both MCAR and MNAR. In both conditions no imputation performs nearly the same or better than the five imputation methods.

Next, we assessed whether imputation can generate additional quantitative peptides. While peptide detection rates have increased significantly over the past decade, not every detected peptide is necessarily quantitative. For a peptide to be considered “quantitative,” increases in measured signal must directly correspond to increases in peptide abundance across a linear range [34]. We obtained data from a serial dilution series experiment (PXD014815) in which peptides were spiked into a background matrix at known concentration. We used a statistical model developed by Pino *et al*. to determine whether each detected peptide was quantitative before and after imputation [34].

The results of this experiment (Figure 4) show that several imputation methods produce new quantitative peptides. MissForest, kNN and NMF each produce large sets of peptides that are quantitative only after imputation (2,768 for MissForest; 1,050 for kNN; 1,128 for NMF). However, MissForest is the only method that increases the *total* number of quantitative peptides relative to no imputation, producing 10,475 quantitative peptides relative to the 7,707 obtained with no imputation.

**Figure 4.**
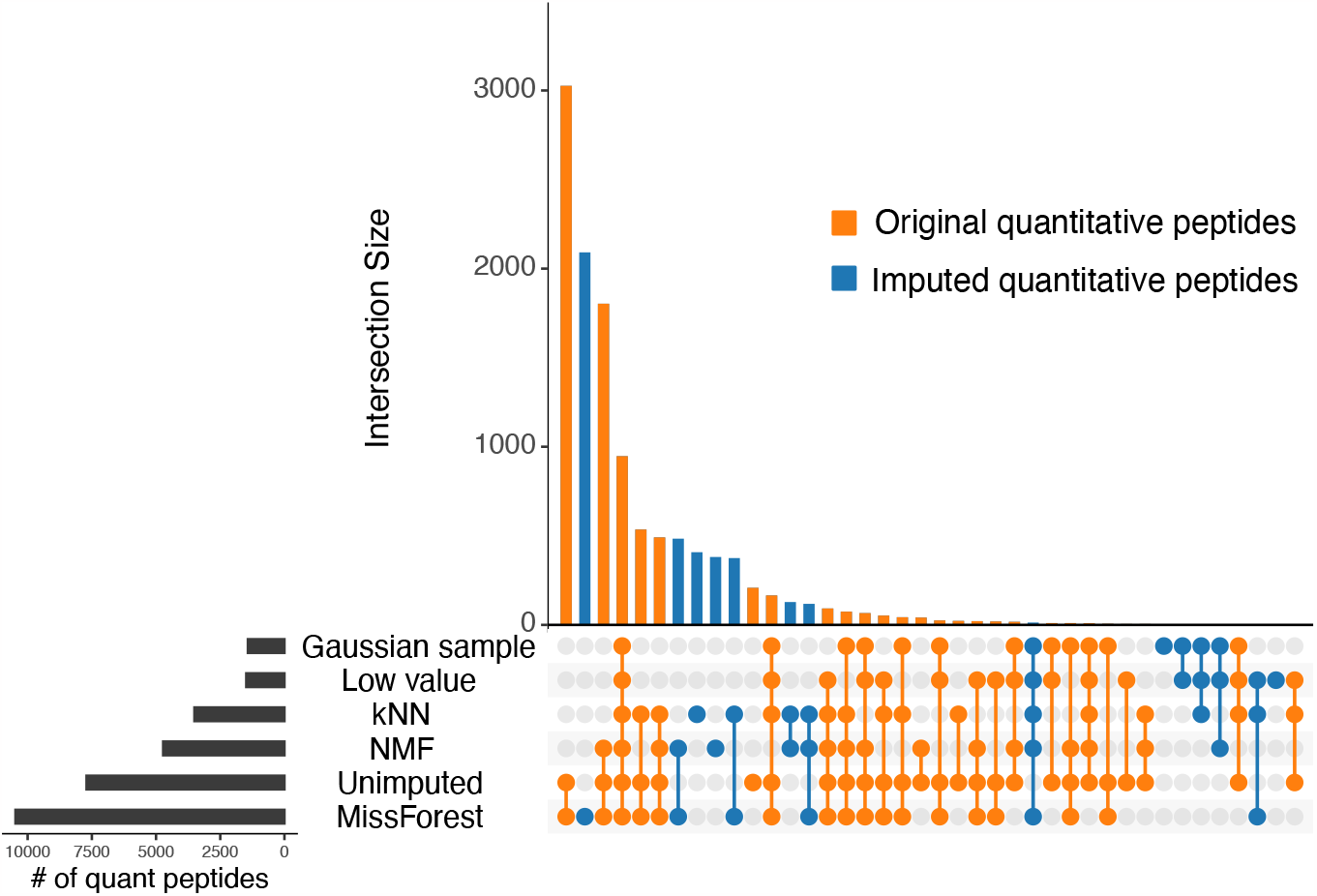
Imputation rescues quantitative peptides. Orange indicates peptides that were quantitative in both the imputed and unimputed data sets. Blue indicates peptides that were only quantitative after imputation. Data was obtained from PXD014815 [34].

We also assessed whether imputation can improve peptide LLOQ, which refers to the minimum abundance at which a peptide can be considered quantitative. For this analysis, we again used the serial dilution data set from Pino *et al*.. We found that while imputation decreases the LLOQ for many peptides, it also increases the LLOQ for many peptides, which is the opposite of the intended effect. Strikingly, MissForest was the only method that decreased the LLOQ of significantly more peptides than it increased (Figure 5, one-sided binomial p-value corrected with Benjamini-Hochberg *<* 0.01).

**Figure 5.**
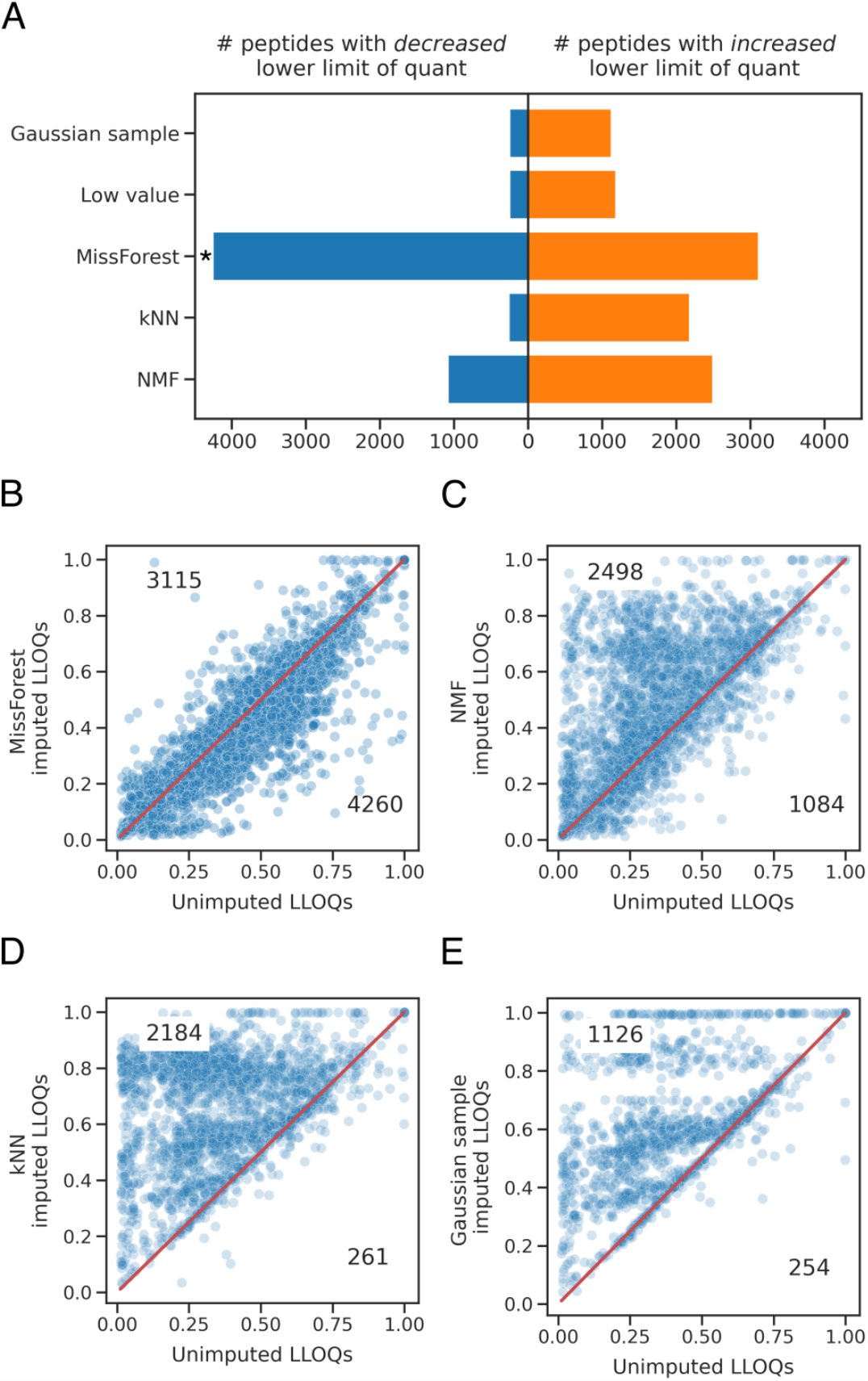
Imputation affects peptide LLOQ. In (A) the asterisk indicates a one-sided binomial Benjamini-Hochberg corrected p-value *<*0.01. In (B–E) the LLOQs of unimputed peptides are plotted against the LLOQs of the same imputed peptides. Only peptides with changes in LLOQ following imputation are plotted. Data was obtained from PXD014815 [34].

For imputation methods to be incorporated into existing proteomics data processing workflows, they must be runnable in a reasonable time frame. We therefore assessed the runtimes of five imputation methods (Figure 6). The two simplest methods, Gaussian sample and low value replacement, typically run in a matter of seconds; NMF and kNN run in a matter of minutes; and MissForest generally takes several hours to complete. Thus, runtime should not present a barrier for incorporation into data processing workflows, with the possible exception of MissForest with its dramatically longer runtime.

**Figure 6.**
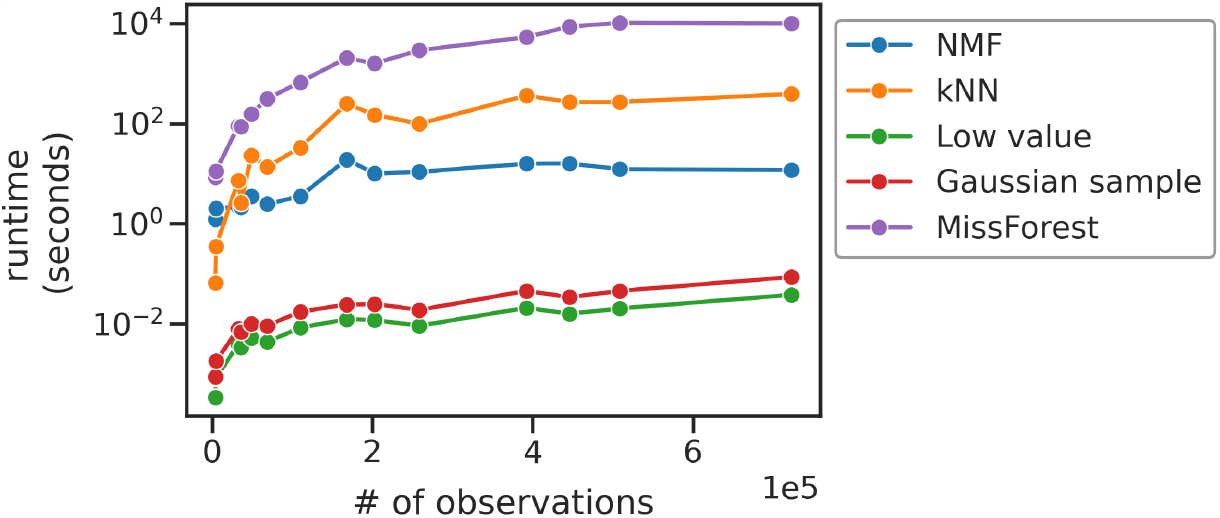
Runtime comparison for imputation methods. Each point represents a public proteomics data set. Data sets are ordered by the number of non-missing observations in their training sets after an 80%/20% MCAR train/test partition. This experiment was performed on a dual CPU Intel Xeon E5-2620 machine with 32 GB RAM. Ten cores were provided for NMF; the remaining methods were run on a single core.

### 3.3 Investigating variance in mass spectrometry-based proteomics data

During the course of this study, we also investigated the extent to which the assumptions underlying several existing imputation approaches are not met by proteomics data. In particular, peptide quantifications are often assumed to be Poisson distributed, and log transformed quantifications are assumed to be Gaussian [22, 23, 50]. One feature of a Poisson distribution is that variance scales linearly with the mean; one feature of a Gaussian distribution is that variance is constant across means. Parametric imputation methods with a Gaussian prior include least-squares regression, the Gaussian sample impute method, and NMF.

To investigate the empirical variance of peptide quantifications, we obtained data from four experiments, each of which contained technical replicates (Table 2). We used three DDA experiments and one DIA. We calculated the means and variances of peptide quantifications across technical replicates, for each detected peptide, for each experiment. We found that peptide quantifications are overdispersed (Figure 7, left); that is, for nearly every peptide, the variance across replicates is greater than the mean intensity across replicates. Log transforming corrected overdispersion for some but not all peptides (Figure 7, right).

**Figure 7.**
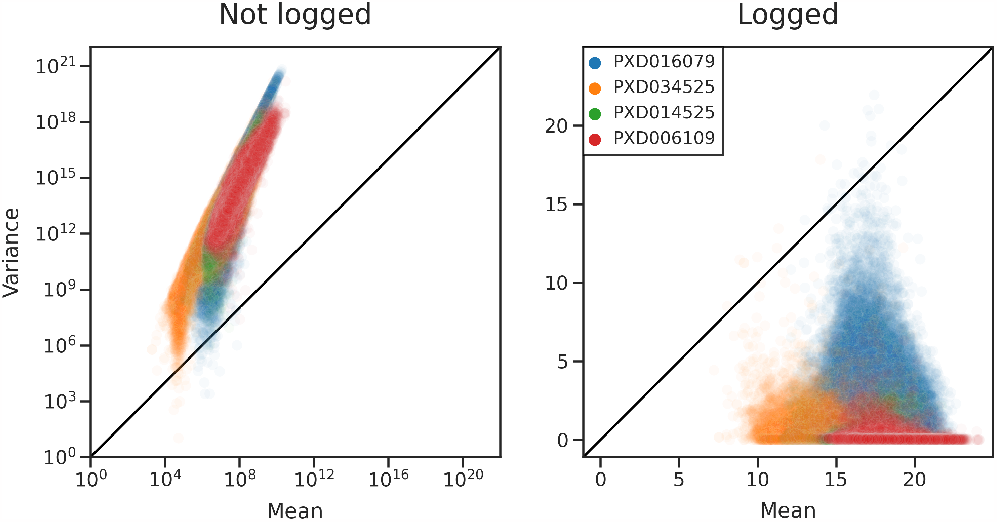
Variance of peptide quantifications is greater than expected. Means and variances were calculated across technical replicates for every detected peptide, for four public proteomics data sets. Each dot corresponds to a peptide. The color scheme is the same in both panels. In the right panel the peptide quantifications have been log transformed.

These results indicate that for two major mass spectrometry acquisition strategies, neither Poisson nor Gaussian assumptions hold. Accordingly, any parametric imputation method with a Gaussian prior is ill-suited for these data. This result also implies that differential expression analysis should be carried out with non-parametric tests such as the Wilcoxon rank sum test, instead of the parametric Student’s t-test.

We also observe that imputation with NMF and MissForest has little effect on the variance of peptide quantifications (Supplementary Figure S3). The Gaussian sample method, however, introduces additional variance. This finding suggests that while NMF and MissForest imputation do not profoundly effect the underlying distribution of peptide quantifications, single-value impute strategies may do so. In this way, single-value impute strategies may introduce artifacts into proteomics data when their underlying assumptions are not met.

## 4 Discussion

The two most popular imputation methods for proteomics data—Gaussian sampling and low value replacement— generally work poorly. This observation is borne out of our evaluation with both traditional and downstream-centric criteria. In the MCAR condition of the traditional evaluation experiment shown in Figure 2, both of these single-value replacement strategies consistently exhibit the worst performance. In the MNAR condition, we expected these two methods to perform well, because both methods assume that missing values are drawn entirely from the low end of the intensity spectrum, an assumption that is met by our MNAR simulation. We indeed see that in the traditional evaluation experiment, the two single-value replacement strategies perform well (Figure 2). However, in the more relevant setting of differential expression imputation, the single-value replacement strategies perform poorly under both MCAR and MNAR conditions (Figure 3). Additionally, neither Gaussian sampling nor low value replacement increase the number of quantitative peptides (Figure 4), and both increase the LLOQ for more peptides than they decrease (Figure 5). These findings suggest that single-value replacement strategies are ill suited to common downstream tasks involving identifying differentially expressed peptides, quantifying low-abundance peptides, or quantifying peptides from small sample volumes. We thus encourage the community to move away from single-value replacement strategies.

Our study also suggests that imputation may not be necessary for differential expression analysis. We find that in the context of identifying ground truth differentially expressed peptides from data with simulated missingness, no imputation works roughly as well as the best imputation methods, in both MCAR and MNAR conditions (Figure 3). In the MCAR condition, the largest AUC value belongs to MissForest at 0.8, only slightly higher than unimputed at 0.76. In the MNAR condition, kNN has the highest AUC at 0.87, and unimputed is close behind with 0.86. Figure 3 reports results for a missingness fraction of 0.25; at missingness fractions of 0.3 and 0.5, unimputed still performs about as well as the best imputation methods (Supplementary Figure S2). In fact, as the missingness fraction increases, unimputed performs better and better relative to the best imputation methods. For example, in the case of MNAR with a 50% missingness fraction, unimputed has an AUC of 0.65, whereas the best imputation method is MissForest with an AUC of 0.55. Taken together, these results cast doubt on the practice of imputing missing values prior to differential expression analysis. This finding is in line with Wolski *et al*., which suggests that statistical models of differential expression that do not impute but rather explicitly model missingness tend to outperform traditional models [51].

We find that imputation can, however, identify new quantitative peptides (Figure 4). The distinction between *quantitative* and *qualitative* peptides is important: as modern proteomic techniques increase the number of identifications, it is worth keeping in mind that not all detected peptides will necessarily be quantitative. We show that MissForest can increase the total number of quantitative peptides in an experiment (Figure 4). Additionally, NMF and kNN can produce new subsets of quantitative peptides, even though they may still decrease the total number of quantitative peptides. Importantly, increasing the number of quantitative peptides will increase the statistical power of any downstream prediction or inference task that relies on peptide abundances. Such tasks include identifying differentially expressed peptides, clustering samples or peptides, identifying co-expression modules, and quality control.

Imputation with MissForest can also improve the LLOQ for a subset of peptides (Figure 5). It is worth acknowledging that while MissForest decreases the LLOQ of significantly more peptides than it increases, it does still increase the LLOQ for a large number of peptides (3,115/24,204 detected peptides). That said, any proteomics study that examines biologically important low-abundance peptides may still benefit from MissForest imputation. As the scale and sensitivity of proteomics experiments increase, MissForest— and future imputation methods—may help researchers study key peptides derived from ever-smaller sample volumes.

Traditional machine learning evaluation criteria do not always agree with downstream-centric criteria. For example, under the MNAR condition of Figure 2, Gaussian sampling and low value replacement both perform exceptionally well, with the lowest reconstruction MSE for three of the six data sets. However, in the context of the dowstream-centric evaluation criteria of Figures 3–5, these two methods consistently perform the worst. This seemingly contradictory finding can once again be reconciled by the fact that single-value impute strategies explicitly assume that missing values are drawn from the low end of the distribution of peptide quantifications; this assumption is met in the MNAR condition of the traditional evaluation experiment. However, the traditional evaluation experiment is somewhat contrived and does not resemble any of the common analysis tasks in modern proteomics. We therefore argue that downstream-centric criteria are more relevant, as they more closely resemble the types of experiments proteomics researchers actually perform. We therefore urge the community to move away from machine learning-style evaluation criteria, as they can mislead with regard to the best performing imputation methods.

Finally, we provide empirical evidence that peptide quantifications exhibit more variance than can be explained under Poisson or Gaussian modeling assumptions (Figure 7). While ion detection may be a Poisson process [22, 23, 50], it is clear that the resulting peptide-level quantifications are not Poisson distributed. One property of a Poisson distribution is that the mean is equal to the variance. We find that this assumption is violated by peptide quantifications: Figure 7 shows that the variance in the intensities of technical replicate peptides is greater than the corresponding means. This result holds true for both DDA and DIA experiments, and it also holds true for protein-level quantifications (Supplementary Figure S4). We speculate that an unaccounted-for noise source is likely present, perhaps electrospray ionization noise. Another assumption is that log-transformed peptide quantifications are roughly Gaussian, that is, variance is constant across all intensity means. We again show that this assumption is violated in both DIA and DDA data: we observe greater-than-expected variance after log transformation.

Future imputation methods should explicitly model the variance present in proteomics data. One obvious choice of generating distribution is the negative binomial distribution, which has an additional parameter that can account for variance independent of the mean. This strategy has been employed previously to model counts from single-cell RNA sequencing experiments [25, 26]. Another option could be to perform variance stabilization prior to imputation. This is the goal with the log transformation; however, as we have shown, logging does not successfully stabilize variance. VSN, a custom variance stabilizing transformation originally developed for microarrays, has been shown to stabilize the variance of protein quantifications [16, 52], as has the generalized log transformation [53]. However, the proteomics community is yet to broadly adopt these methods. Proteomics may also benefit from the variance stabilization technique developed by Bayat *et al*., in which a variance stabilizing function is empirically learned from the data [24]. Successful modeling and variance stabilization approaches could benefit not just imputation but general data denoising procedures for quantitative proteomics.

We speculate that the unusual dimensionality of peptide-by-sample matrices, generally thousands of peptides by less than 100 samples, may cause problems for existing imputation methods, many of which were originally developed for microarray imputation and relatively square matrices. Future imputation methods may benefit from explicitly accounting for unusual dimensionality of these matrices.

The proteomics community would benefit from easy-to-use and broadly applicable imputation methods. As previously reported [1, 2, 14, 15], we find that the best choice in imputation method depends on the analysis task and the details of the experiment. For certain tasks, existing imputation methods may not be useful at all. This suggests the need for new imputation methods that are generalizable enough to accurately handle data from any acquisition strategy and type of missingness. Deep neural networks have proven highly generalizable in other contexts. Recent “deep” impute methods may be a step in the right direction [20], though much work remains to be done. In the future, data-driven imputation methods will likely be broadly adopted as part of general signal processing workflows in proteomics.

## Associated Content

The code used to generate all the figures in this manuscript, as well as the Supplementary Material, can be found at: https://github.com/Noble-Lab/2023-prot-impute-benchmark. The custom NMF imputation model can be found at https://github.com/Noble-Lab/ms_imputer. All data sets used in this study are publicly available and can be found on PRIDE, CPTAC or Panorama [27, 29, 54].

## References

[1] Bramer L, Irvahn J, Piehowski P, Rodland K, and Webb-Robertson BJ. A review of imputation strategies for isobaric labeling-based shotgun proteomics. Journal of Proteome Research, 20:1–13, 2021.

[2] Webb-Robertson BJ, Wiberg H, Matzke M, Brown J, Wang J, McDermott J, Smith R, Rodland K, Metz T, Pounds J, and Waters K. Review, evaluation and discussion of challenges of missing value imputation for mass spectrometry-based label-free global proteomics. Journal of Proteome Research, 14:1993–2001, 2015.

[3] Troyanskaya O, Cantor M, Sherlock G, Brown P, Hastie T, Tibshirani R, Botstein D, and Altman R. Missing value estimation method for DNA microarrays. Bioinformatics, 17:520–525, 2001.

[4] Sterne J, White I, Carlin J, Spratt M, Royston P, Kenward M, Wood A, and Carpenter J. Multiple imputation for missing data in epidemiological and clinical research: potential and pitfalls. BMJ, 338(b2393), 2009.

[5] Keerin P and Boongoen T. Estimation of missing values in astronomical survey data: An improved local approach using cluster directed neighbor selection. Information Processing and Management, 59(102881), 2022.

[6] Luken K, Padhy R, and Wang XR. Missing data imputation for galaxy redshift estimation. NeurIPS, 2021.

[7] Linderman G, Zhao J, Roulis M, Bielecki P, Flavell R, Nadler B, and Kluger Y. Zero-preserving imputation of single-cell RNA-seq data. Nature Communications, 192, 2022.

[8] van Dijk D, Sharma R, Nainys J, Yim K, Kathail P, Carr A, Burdziak C, Moon K, Chaffer C, Pattabi- raman D, Bierie B, Mazutis L, Wolf G, Krishnaswamy S, and Pe’er D. Recovering gene interactions from single-cell data using data diffusion. Cell, 174:716–729, 2018.

[9] Tyanova S, Temu T, Sinitcyn P, Carlson A, Hein M, Geiger T, Mann M, and Cox J. The Perseus computational platform for comprehensive analysis of (prote)omics data. Nature Methods, 13:731–740, 2016.

[10] Kowarik A and Templ M. Imputation with the R package VIM. Journal of Statistical Software, 74(7), 2016.

[11] Stekhoven D and Buhlmann P. MissForest –non-parametric missing value imputation for mixed-type data. Bioinformatics, 28:112–118, 2012.

[12] Stacklies W, Redestig H, Scholz M, Walther D, and Selbig J. pcaMethods—a bioconductor package providing PCA methods for incomplete data. Bioinformatics, 23(9), 2007.

[13] Josse J and Husson F. missMDA: a package for handling missing values in multivariate data analysis. Journal of Statistical Software, 70(1), 2016.

[14] Egert J, Brombacher E, Warscheid B, and Kreutz C. DIMA: Data-driven selection of an imputation algorithm. Journal of Proteome Research, 20:3489–3496, 2021.

[15] Lazar C, Gatto L, Ferro M, Bruley C, and Burger T. Accounting for the Multiple Natures of Missing Values in Label-Free Quantitative Proteomics Data Sets to Compare Imputation Strategies. Journal of Proteome Research, 15:1116–1125, 2016.

[16] Välikangas T, Suomi T, and Elo L. A comprehensive evaluation of popular proteomics software workflows for label-free proteome quantification and imputation. Briefings in Bioinformatics, 19(6), 2018.

[17] Dabke K, Kreimer S, Jones M, and Parker S. A simple optimization workflow to enable precise and accurate imputation of missing values in proteomic data sets. Journal of Proteome Research, 20:3214–3229, 2021.

[18] Xu J, Wang Y, Xu X, Cheng KK, Raftery D, and Dong J. NMF-Based Approach for Missing Values Imputation of Mass Spectrometry Metabolomics Data. Molecules, 26(19), 2021.

[19] Hediyeh Zadeh S, Webb A, and Davis M. MSImpute: Imputation of label-free mass spectrometry peptides by low-rank approximation. bioRxiv, 2020.

[20] Webel H, Niu L, Nielsen AB, Locard-Paulet M, Mann M, Jensen LJ, and Rasmussen S. Mass spectrometry-based proteomics imputation using self supervised deep learning. bioRxiv, 2023.

[21] Plubell D, Kall L, Webb-Robertson BJ, Bramer L, Ives A, Kelleher N, Smith L, Montine T, Wu C, and MacCoss M. Putting Humpty Dumpty Back Together Again: What Does Protein Quantification Mean in Bottom-Up Proteomics? Journal of Proteome Research, 21:891–898, 2022.

[22] Ipsen A. Derivation from first principles of the statistical distribution of the mass peak intensities of MS data. Analytical Chemistry, 87:1726–1734, 2015.

[23] Ipsen A and Ebbels T. Prospects for a statistical theory of LC/TOFMS data. Journal of the American Society of Mass Spectrometry, 23:779–791, 2012.

[24] Bayat F and Libbrecht M. VSS: variance-stabilized signals for sequencing-based genomic signals. Bioinformatics, 3723), 2021.

[25] Risso D, Perraudeau F, Gribkova S, Dudoit S, and Vert JP. A general and flexible method for signal extraction from single-cell RNA-seq data. Nature Communications, 9(284), 2018.

[26] Hafemeister C and Satija R. Normalization and variance stabilization of single-cell RNA-seq data using regularized negative binomial regression. Genome Biology, 20(296), 2019.

[27] Perez-Riverol Y, Csordas A, Bai J, Bernal-Llinares M, Hewapathirana S, Kundu D, Inuganti A Griss J, Mayer G, Eisenacher M, Perez E, Uszkoreit J, Pfeuffer J, Sachsenberg T, Yilmaz S, Tiwary S, Cox J, Audain E, Walzer M, Jarnuczak A, Ternent T, Brazma A, and Vizcaino JA. The PRIDE database and related tools and resources in 2019: improving support for quantification data. Nucleic Acids Research, 8:442–450, 2019.

[28] Vizcaíno J, Deutsch E, Wang R, Csordas A, Reisinger F, Ríos D, Dianes J, Sun Z, Farrah T, Bandeira N, Binz PA, Xenarios I, Eisenacher M, Mayer G, Gatto L, Campos A, Chalkley R, Kraus HJ, Albar JP, Martinez-Bartolomé S, Apweiler R, Omenn G, Martens L, Jones A, and Hermjakob H. ProteomeXchange provides globally coordinated proteomics data submission and dissemination. Nature Biotechnology, 32:223–226, 2014.

[29] Edwards N, Oberti M, Thangudu R, Cai S, McGarvey P, Jacob S, Madhavan S, and Ketchum K. The CPTAC Data Portal: A Resource for Cancer Proteomics Research. Journal of Proteome Research, 14:2707–2713, 2015.

[30] Selamoglu N, Önder Ö, Öztürk Y, Khalfaoui-Hassani B, Blaby-Hass C, Garcia B, Koch HG, and Daldal F. Comparative differential cuproproteomes of Rhodobacter capsulatus reveal novel copper homeostasis related proteins. Metallomics, 12(572), 2020.

[31] Meier F, Geyer P, Virreira Winter S, Cox J, and Mann M. BoxCar acquisition method enables single-shot proteomics at a depth of 10,000 proteins in 100 minutes. Nature Methods, 15:440–448, 2018.

[32] Bekker-Jensen D, Bernhardt O, Hogrebe A, Martinez-Val A, Verbeke L, Gandhi T, Kelstrup C, Reiter L, and Olsen J. Rapid and site-specific deep phosphoproteome profiling by data-independent acquisition without the need for spectral libraries. Nature Communications, 11(787), 2020.

[33] Merrihew G, Park J, Plubell D, Searle B, Keene D, Larsen E, Bateman R, Perrin R, Chhatwal J, Farlow M, McLean C, Ghetti B, Newell K, Frosch M, Montine T, and MacCoss M. A peptide-centric quantitative proteomics dataset for the phenotypic assessment of Alzheimer’s disease. bioRxiv, 2022.

[34] Pino L, Searle B, Yang HY, Hoofnagle A, Noble W, and MacCoss M. Matrix-matched calibration curves for assessing analytical figures of merit in quantitative proteomics. Journal of Proteome Research, 19:1147–1153, 2020.

[35] Nitschko V, Kunzelmann S, Frohlich T, Arnold G, and Forstemann K. Trafficking of siRNA precursors by the dsRBD protein blanks in Drosophila. Nucleic Acids Research, 48(7), 2020.

[36] Azizan A, Kaschani F, Barinas H, Blaskowski S, Kaiser M, and Denecke M. Using proteomics for an insight into the performance of activated sludge in a lab-scale WWTP. International Biodeterioration and Biodegradation, 149(104934), 2020.

[37] Murugaiyan J, Eravci M, Weise C, Roesler U, Sprague L, Neubauer H, and Wareth G. Pan-proteomic analysis and elucidation of protein abundance among the closely related Brucella species, Brucella abortus and Brucella melitensis. Biomolecules, 10(836), 2020.

[38] Rodrigues D, Mufteev M, Weatheritt R, Djuric U, Ha K, Ross PJ, Wei W, Piekna A, Sartori M, Byres L, Mok R, Zaslavsky K, Pasceri P, Diamandis P, Morris Q, Blencowe B, and Ellis J. Shifts in ribosomal engagement impact key gene sets in neurodevelopment and ubiquitination in Rett syndrome. Cell Reports, 30:4179–4196, 2020.

[39] Petralia F, Tignor N, Reva B, Koptyra M, Chowdhury S, Rykunov D, Krek A, Ma W, Zhu Y, Ji J, Calinawan A, Whiteaker J, Colaprico A, Stathias V, Omelchenko T, Song X, Raman P, Guo Y, Brown M, Ivey R, Szpyt J, Thakurta SG, Gritsenko M, Weitz K, Lopez G, Kalayci S, Gumus Z, Yoo S, da Veiga Leprevost F, Chang HY, Krug K, Katsnelson L, Wang Y, Kennedy J, Voytovich U, Zhao L, Gaonkar K, Ennis B, Zhang B, Baubet V, Tauhid L, Lilly J, Mason J, Farrow B, Young N, Leary S, Moon J, Petyuk V, Nazarian J, Adappa N, Palmer J, Lober R, Rivero-Hinojosa S, Wang LB, Wang J, Broberg M, Chu R, Moore R, Monroe M, Zhao R, Smith R, Zhu J, Robles A, Mesri M, Boja E, Hiltke T, Rodriguez H, Zhang B, Schadt E, Mani DR, Ding L, Iavarone A, Wiznerowicz M, Schurer S, Chen XS, Heath A, Rokita JL, Nesvizhskii A, Fenyo D, Rodland K, Liu T, Gygi S, Paulovich A, Resnick A, Storm P, Roo B, Wang P, Children’s Brain Tumor Network, and Clinical Proteomic Tumor Analysis Consortium. Integrated proteogenomic characterization across major histological types of pediatric brain cancer. Cell, 183:1962–1985, 2020.

[40] Satpathy S, Jaehnig E, Krug K, Kim BJ, Saltzman A, Chan D, Holloway K, Anurag M, Huang C, Singh P, Gao A, Namai N, Dou Y, Wen B, Vasaikar S, Mutch D, Watson M, Ma C, Ademuyiwa F, Rimawi M, Schiff R, Hoog J, Jacobs S, Malovannaya A Hyslop T, Clauser K, Mani D, Perou C, Miles G, Zhang B, Gillette M, Carr S, and Ellis M. Microscaled proteogenomic methods for precision oncology. Nature Communications, 11(532), 2020.

[41] O’Connell J, Paulo J, O’Brien J, and Gygi S. Proteome-wide evaluation of two common protein quantification methods. Journal of Proteome Research, 17(5), 2018.

[42] Cox J and Mann M. MaxQuant enables high peptide identification rates, individualized p.p.b.-range mass accuracies and proteome-wide protein quantification. Nature Biotechnology, 26:1367–1372, 2008.

[43] Searle B, Pino L, Egertson J, Ting Y, Lawrence R, MacLean B, Villen J, and MacCoss M. Chromatogram libraries improve peptide detection and quantification by data independent acquisition mass spectrometry. Nature Communications, 9(5128), 2018.

[44] MacLean B, Tomazela D, Shulman N, Chambers M, Finney G, Frewen B, Kern R, Tabb D, Liebler D, and MacCoss M. Skyline: an open source document editor for creating and analyzing targeted proteomics experiments. Bioinformatics, 26(7), 2010.

[45] da Veiga Leprevost F, Haynes S, Avtonomov D, Chang HY, Shanmugam A, Mellacheruvu D, Kong A, and Nesvizhskii A. Philosopher: a versatile toolkit for shotgun proteomics data analysis. Nature Methods, 17:869–870, 2020.

[46] Benjamini Y and Hochberg Y. Controlling the false discovery rate: a practical and powerful approach to multiple testing. Journal of the Royal Statistical Society, 57(1), 1995.

[47] Conway J, Lex A, and Gehlenborg N. UpSetR: an R package for the visualization of intersecting sets and their properties. Bioinformatics, 33(18), 2017.

[48] Andrews T and Hemberg M. False signals induced by single-cell imputation. F1000 Research, 7(1740), 2019.

[49] Ly LH and Vingron M. Effect of imputation on gene network reconstruction from single-cell RNA-seq data. Patterns, 3(100414), 2022.

[50] Kimmel J, Kyu Yoon O, Zuleta I, Trapp O, and Zare R. Peak height precision in Hadamard transform time-of-flight mass spectra. American Society of Mass Spectrometry, 16(1117-1130), 2005.

[51] Wolski W, Nanni P, Grossmann J, d’Errico M, Schlapbach R, and Panse C. prolfqua: A comprehensive R-package for proteomics differential expression analysis. Journal of Proteome Research, 2023.

[52] Huber W, von Heydebreck A, Sultmann H, Poustka A, and Vingron M. Variance stabilization applied to microarray data calibration and to the quantification of differential expression. Bioinformatics, 18(Supp 1), 2002.

[53] Anderle M, Roy S, Lin H, Becker C, and Joho K. Quantifying reproducibility for differential proteomics: noise analysis for protein liquid chromatography-mass spectrometry of human serum. Bioinformatics, 20(18), 2004.

[54] Sharma V, Eckels J, Schilling B, Ludwig C, Jaffe J, MacCoss M, and MacLean B. Panorama Public: A Public Repository for Quantitative Data Sets Processed in Skyline. Molecular and Cellular Proteomics, 17(6), 2018.

