## Supplement for "Evaluating proteomics imputation methods with improved criteria"

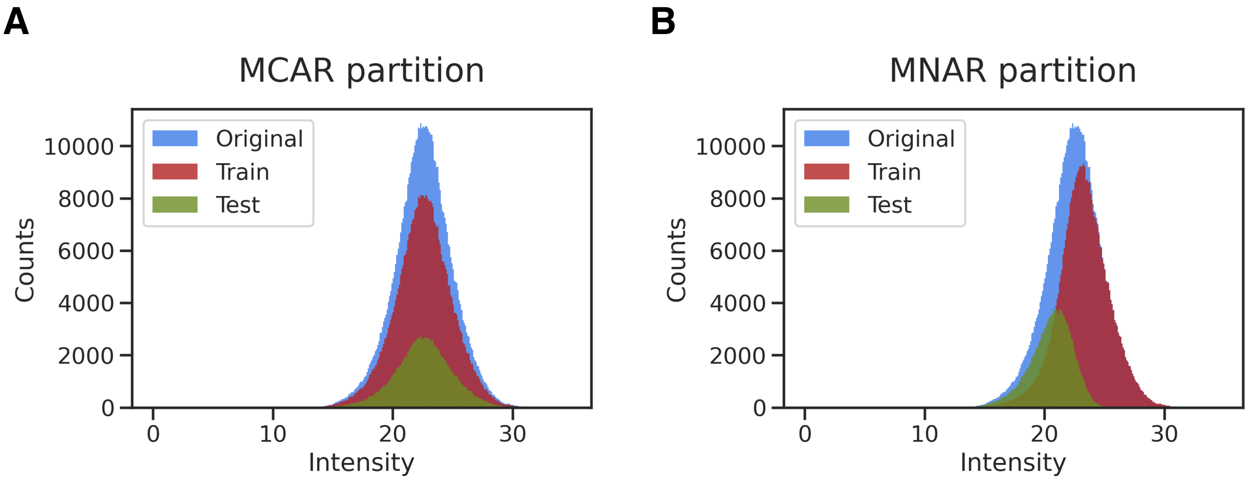


Supplementary Figure S1. **Distributions of partitions for both MCAR and MNAR.** Data from PXD03452 [33]. Log-transformed peptide-level quantifications are shown. In each case, 25% of present matrix entries were selected for the test set.

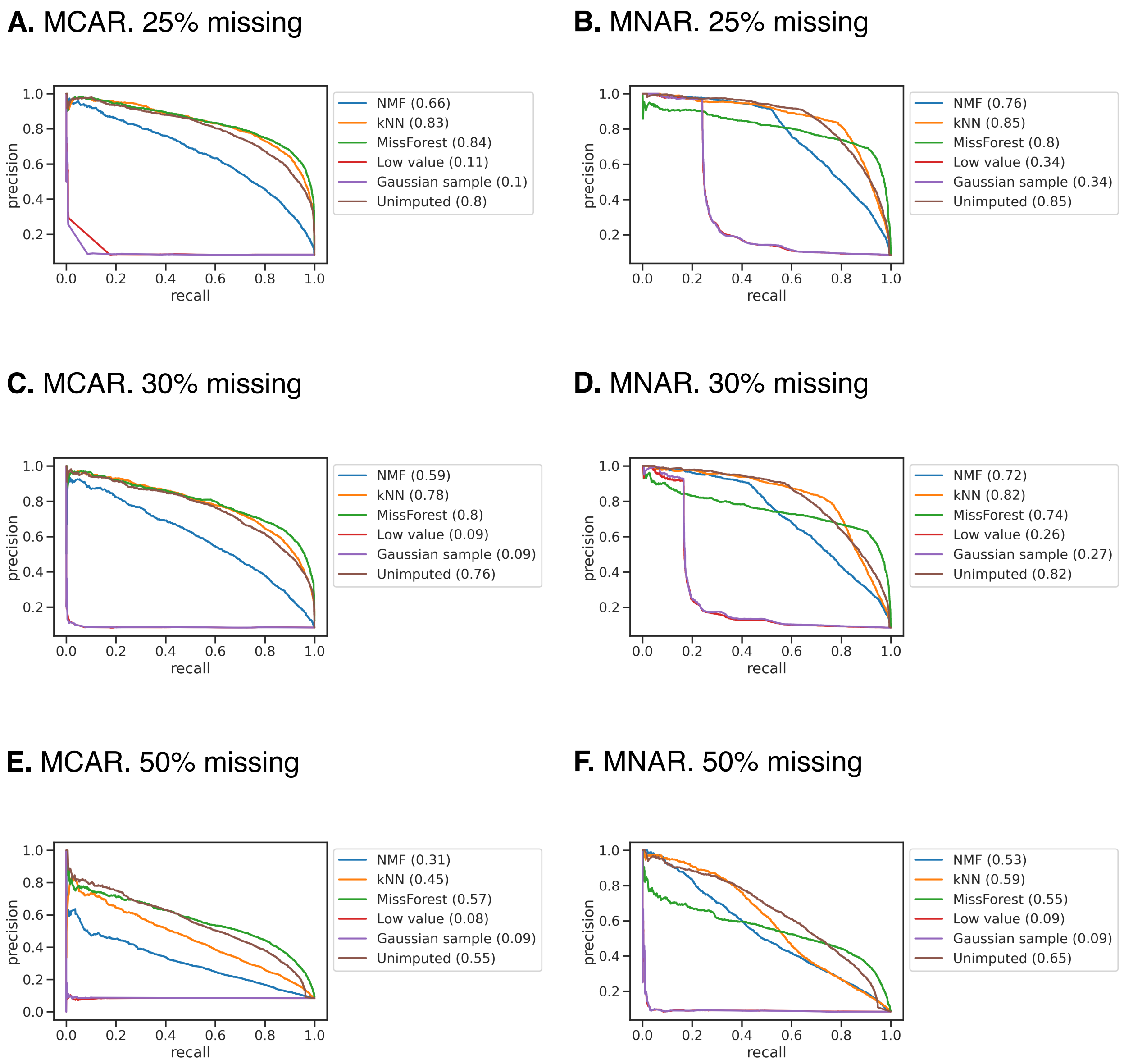


Supplementary Figure S2. **Additional evaluation of the ability of imputation methods to reconstruct differentially expressed peptides.** Data obtained from PXD03452, as in Figure 3 [33]. Differentially expressed peptides were identified between two experimental groups for Alzheimer's disease DIA data. AUC values are shown in parentheses.

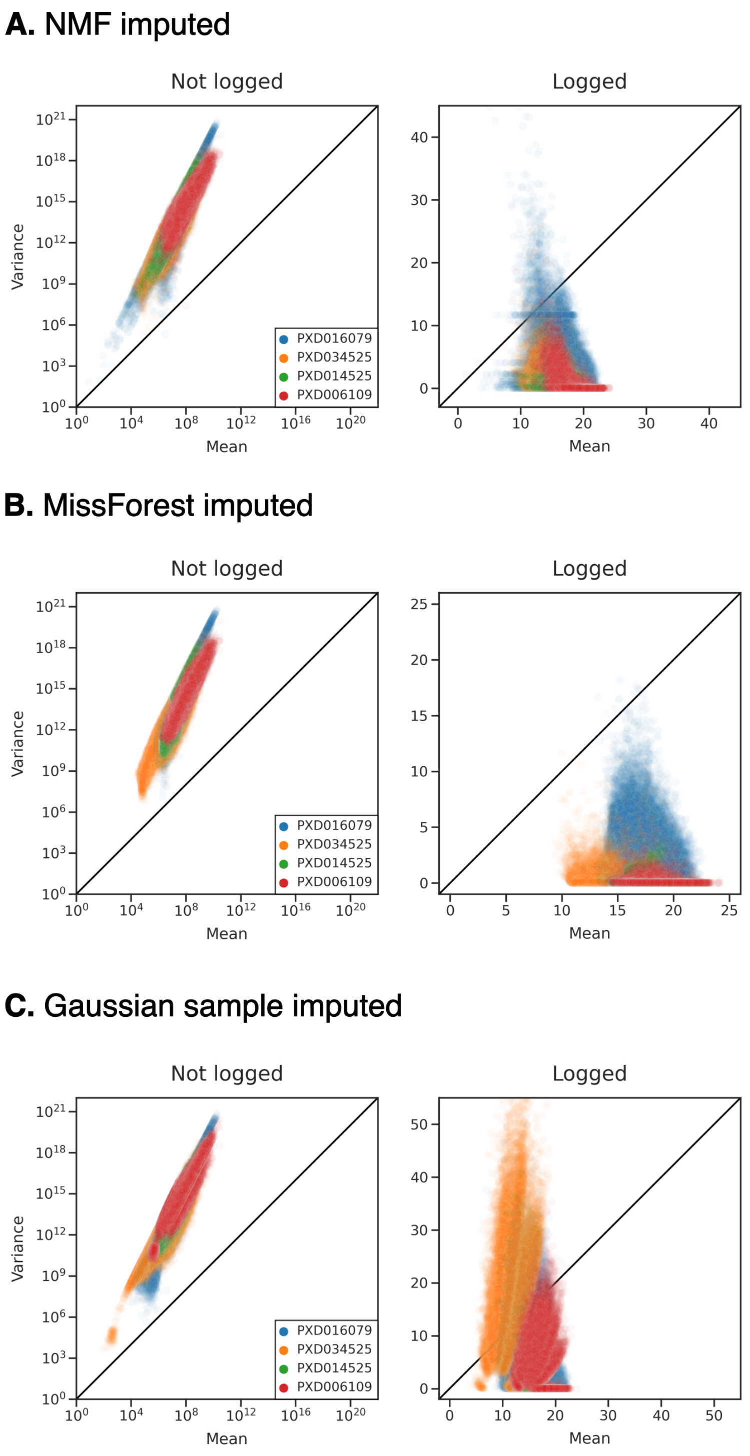


Supplementary Figure S3. **Variance of peptide quantifications is greater than expected following imputation.** Four public proteomics data sets were imputed with NMF, MissForest and Gaussian sampling. Means and variances were calculated across technical replicates for each imputed peptide, for each data set. In the righthand panels, imputed quantifications have been log-transformed.

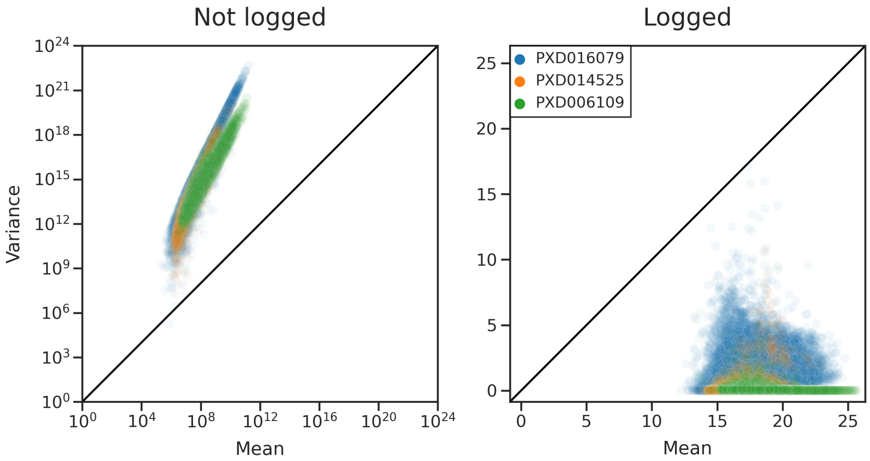


Supplementary Figure S4. **Variance of *protein* quantifications is greater than expected.** Means and variances were calculated across technical replicates for each protein, for three public protein-level quantification data sets. In the righthand panel protein quantifications have been log-transformed. Color scheme is the same in both panels.
